# Why not both? Measuring cortisol and corticosterone in poison frogs

**DOI:** 10.1101/2023.06.19.545597

**Authors:** Sarah E. Westrick, Ryan T. Paitz, Eva K. Fischer

## Abstract

A general tenet in stress physiology is that the hypothalamic-pituitary-adrenal (HPA) axis predominantly produces one glucocorticoid (GC) in response to stressors. However, most vertebrates produce both cortisol and corticosterone, and these steroids show variation across species in absolute levels, relative proportions, and stress responsivity and regulate much more than just stress physiology. Therefore, focusing on a single GC within a species may not capture the whole story. In the present study, we set out to validate non-invasive waterborne hormone measurements in our focal species, the dyeing poison frog *Dendrobates tinctorius*. In pursuing this goal, we uncovered unexpected patterns of GC abundance within and across species of poison frogs. *D. tinctorius* had higher amounts of corticosterone than cortisol in both plasma and waterborne samples, and corticosterone was responsive to adrenocorticotropic hormone (ACTH) as canonically assumed. However, corticosterone and cortisol levels were surprisingly similar in *D. tinctorius*, and cortisol was more abundant than corticosterone in water samples from four additional poison frog species. These results challenge the broadly accepted assumption that corticosterone is always more abundant in amphibians and add to the growing literature highlighting the importance of measuring both GCs to understand (stress) physiology.

## 1. Introduction

The ability to measure physiology non-invasively in the lab and in the wild is key for behavioral ecologists and conservation biologists seeking to understand animal behavior in a changing world. Glucocorticoids – most commonly cortisol and corticosterone produced by the adrenal or interrenal glands (Sapolsky et al., 2000) – are of keen interest because they elicit a variety of physiological, morphological, and behavioral responses (MacDougall-Shackleton et al., 2019). Glucocorticoids are often measured from plasma samples but can also be measured in feces, urine, hair, feathers, baleen, eggs, and water (Palme, 2019). These non-invasive sampling methods can reduce the confounds of stress from sampling itself and allow for repeated sampling from small individuals where repeated blood draws are not possible or when individuals cannot be captured (Palme, 2019). The majority of studies measure only a single GC, the one considered to be “dominant” and a readout of stress (Hancock, 2010). Which GC is dominant varies across species (Koren et al., 2012), with the general assumption that this variation is not functionally interesting because the two GCs are physiologically redundant. Yet a key thing that is often overlooked in stress physiology is the fact that other glucocorticoids are present, potentially in substantial amounts, and can be produced independently of the HPA/I axis (Bornstein et al., 2008), for example locally in different tissue types (Schmidt and Soma, 2008). Moreover, there is intriguing evidence that GCs long assumed to be physiologically interchangeable may in fact play distinct, specialized roles (e.g. Schmidt and Soma, 2008). To fully understand the complex roles glucocorticoids play, we need to move beyond studying a single GC to characterization of multiple GCs that together regulate behavior, development, and individual and population health. In the present study, we set out to validate non-invasive waterborne hormone measurements in our focal species, the dyeing poison frog *Dendrobates tinctorius*. In pursuing this goal, we uncovered unexpected patterns of cortisol and corticosterone abundance within and across species of poison frogs, with implications for how we think about and quantify GC physiology. Below, we briefly review the literature comparing cortisol and corticosterone to provide context for our study and its implications. Our findings highlight the importance of this question specifically in frogs and for those interested in behavior and conservation across vertebrates more broadly.

### 1.1 Cortisol vs corticosterone

Cortisol and corticosterone are best known for their role in the “stress response” during which they coordinate behavioral, physiological, and metabolic responses. However, they also change as a result of ACTH-dependent circannual and circadian rhythms (Dickmeis et al., 2013) and play a critical role in other physiological processes (MacDougall-Shackleton et al., 2019), including, but not limited to, energy metabolism (Vegiopoulos and Herzig, 2007), immune response (Cruz-Topete and Cidlowski, 2015), growth and development (Seckl and Meaney, 2004), reproduction (Whirledge and Cidlowski, 2017), and cardiovascular function (Cruz-Topete et al., 2016). Circulating GCs also impact behaviors including aggression (Haller, 2014), parental care (Bales et al., 2006; Crossin et al., 2012; Raulo and Dantzer, 2018), and feeding/foraging (Crespi and Denver, 2005; Crossin et al., 2012) to name a few. Because of their widespread influences, changes in GC levels and/or their receptors can be immensely impactful on survival and fitness. While acutely elevated GC levels often have adaptive functions, chronically elevated GCs disrupt normal physiology and decrease fitness (reviewed in Bonier et al., 2009 and Breuner et al., 2008).

Many vertebrates produce both cortisol and corticosterone. As steroid hormones, cortisol and corticosterone are produced from cholesterol via partially overlapping biosynthetic pathways (Fig. 1A) (Payne and Hales, 2004). While these pathways share precursors, the final synthesis steps are non-overlapping, and the two GCs cannot be directly converted from one to the other (Fig. 1A). Typically, either cortisol or corticosterone is considered the dominant GC, defined as the one released in response to adrenocorticotropin hormone (ACTH). Therefore, ACTH challenges are commonly used to measure the responsiveness of the adrenals, to identify the dominant GC, and to validate that a chosen method of hormone measurement is detecting an increase in HPA axis activity (Palme, 2019; Touma and Palme, 2005). When ACTH challenges are not possible, dominance is assumed based on higher abundance (Aerts, 2018), resulting in conflation of dominance and abundance. Because both cortisol and corticosterone bind to glucocorticoid and mineralocorticoid receptors, they are often assumed to be entirely interchangeable in their physiological function (e.g. Funder, 1993).

**Figure 1.**
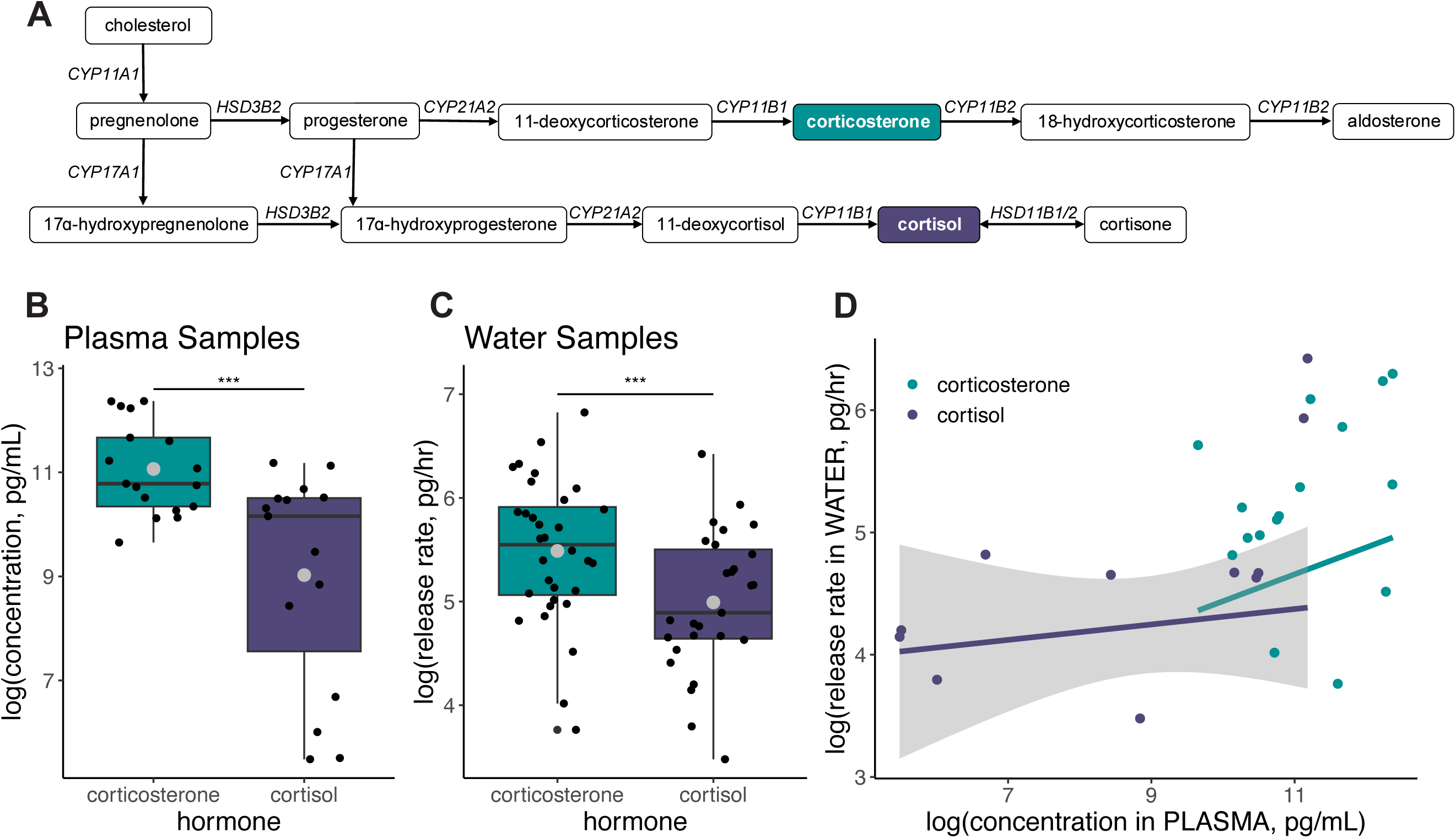
(A) Cortisol and corticosterone are produced through similar pathways which share biosynthetic enzymes. However, the pathway to produce cortisol requires the hydroxylase enzyme cytochrome P450 17A1, encoded by the gene *CYP17A1,* to convert pregnenolone to 17ɑ-hydroxypregnenolone or convert progesterone to 17ɑ-hydroxyprogesterone. Notably, corticosterone is not converted directly into cortisol or vice versa. Corticosterone was more abundant in both (B) plasma and (C) water samples from *D. tinctorius*. Purple boxplots indicate cortisol data. Boxplots for corticosterone data are in teal. Grey dots indicate the mean for each treatment group. Black dots are individual water samples. (*** = p < 0.001). (D) The concentration of cortisol, but not corticosterone, in plasma was positively correlated with the concentration in water samples from the same individuals. Grey shaded area is the confidence interval around the statistically significant linear regression for cortisol.

Due to taxonomic patterns and broadly accepted assumptions of physiological interchangeability, we often make assumptions about which GC is dominant, and therefore assumed to be more abundant, in our species of interest. Consequently, many studies measure only one GC, such as cortisol in fish or corticosterone in birds. These simplifying assumptions are especially prevalent in studies using GCs as a readout of stress or correlate of behavior, rather than exploring GC physiology per se. Yet, these assumptions and generalizations can be misleading.

GC dominance and the ratio of corticosterone to cortisol varies both across (Koren et al., 2012) and within species (e.g., Keogh et al., 2020; Koren et al., 2012; Larson et al., 2009), and a growing number of studies demonstrate the importance of testing interspecific variation in GCs even among closely related species. For instance, across mammals, we see all combinations of which GC responds to an acute stressor or exogenous ACTH, independent of relative abundance. Some mammals respond primarily with an increase in cortisol (Ganjam et al., 1972; Johnston et al., 2013; Van Mourik et al., 1985; Vera et al., 2012, 2011), while in others corticosterone increases (Young et al., 2004), and still in others *both* GCs respond (Rosenthal et al., 1993; Smith and Bubenik, 1990; Young et al., 2004). Sea otters (*Enhydra lutris*) are an interesting case where some studies document an increase in cortisol in response to ACTH (Murray et al., 2020; Wasser et al., 2000), while others find an acute stressor (catching and handling) causes an increase in corticosterone (Larson et al., 2009). These findings suggest a potential for non-ACTH regulation of corticosterone, such as modulation by epinephrine or norepinephrine (Bornstein and Chrousos, 1999). This within and among species variance in abundance and responsiveness suggests that we should use caution when following traditional assumptions about which GC to measure based on order, family, or even population within the same species (Hancock, 2010).

More importantly, the GC that is less abundant or not responsive to ACTH can still play an important physiological role. For instance, in humans, cortisol is considered the primary effector hormone of the HPA axis (Brück, 1983); however, corticosterone plays a role in the central nervous system (Raubenheimer et al., 2006) and pre-natal responses to stressors (Wynne-Edwards et al., 2013). Corticosterone is also an intermediary in the production of aldosterone and the rate of conversion from corticosterone to aldosterone may vary resulting in more or less circulating corticosterone in cortisol-dominant species (Koren et al., 2012). Additionally, steroid receptors and transporters can interact with cortisol and corticosterone differently. In fact, the two GCs can elicit different effects due to different receptor binding affinities (Sutanto and De Kloet, 1987). For example, in mice, P-glycoprotein transports cortisol, but not corticosterone, out of the brain (Karssen et al., 2001). By measuring only one GC, we miss out on finding species that produce measurable amounts of both, miss opportunities to test hypotheses about the independent regulation and function of cortisol vs. corticosterone, and do not have a complete picture of physiological regulation in health and disease.

### 1.2 Glucocorticoids in amphibians

Environmental and anthropogenic changes are contributing to amphibian population decline and rapid biodiversity loss. These stressors include global climate change, pesticides, pollution, and habitat loss or alteration (Blaustein and Kiesecker, 2002; Collins, 2010; Kiesecker, 2011; Kiesecker et al., 2001; Ohmer et al., 2021; Stuart et al., 2004). Many amphibians are small bodied and repeated plasma sampling is not feasible. Developing non-invasive techniques to study the stress physiology of amphibians is instrumental in understanding how species respond to multifactor environmental stressors, such as habitat modification, climate variability, and disease, and can inform conservation strategies for at-risk species (Kumar and Umapathy, 2019; Narayan, 2013; Palme, 2019; Walls and Gabor, 2019). GCs also play critical roles in metamorphosis in amphibians (Denver, 2009; Kulkarni and Buchholz, 2014) with size at metamorphosis having life-long consequences for survival and fitness (Glennemeier and Denver, 2002; Kulkarni and Gramapurohit, 2017).

Corticosterone is typically referenced as the dominant and vastly more abundant glucocorticoid in amphibians (Moore and Jessop, 2003; Norris and Carr, 2013; Sandor, 1969; Wingfield and Romero, 2011), and occasionally listed as amphibians’ only glucocorticoid (Katsu and Iguchi, 2016). In the 1970s and 1980s, early studies of steroid synthesis reported production of corticosterone and little, if any, cortisol from isolated interrenal cells of frogs and toads *in vitro* (Chan and Edwards, 1970; Jolivet-Jaudet and Ishii, 1983; Jolivet-Jaudet and Leloup-Hatey, 1984; Mehdi and Carballeira, 1971). The results of these studies from just a few species of frogs and toads (namely *Bufo bufo, Rana catesbeiana*, and *Xenopus laevis*) led to a strong focus on corticosterone in most studies on stress physiology across all amphibians.

This bias towards studying corticosterone in amphibians prevails despite a history of cortisol being detected in several species of anurans (frogs and toads) (e.g., Dale, 1962; Krug et al., 1983) and a growing number of studies demonstrating that the assumption of corticosterone responsiveness to ACTH/acute stressors and/or greater corticosterone abundance does not hold for all amphibians. For instance, in fecal samples from two species of Panamanian harlequin frogs (*Atelopus certus* and *Atelopus glyphus*) both cortisol and corticosterone were responsive to ACTH, but cortisol metabolites were detected at much higher concentrations overall (Cikanek et al., 2014). In the common water frog (*Rana esculenta*), cortisol and corticosterone are both detectable in plasma and both are elevated in ovulatory and postovulatory phases compared to the preovulatory phase (Gobbetti and Zerani, 1993).

In an attempt to resolve these discrepancies, some researchers have suggested aquatic amphibia may rely on cortisol as their primary GC, as many fishes do, whereas semi-terrestrial or terrestrial amphibia use corticosterone (Jungreis et al., 1970). For example, aquatic hellbender salamanders (*Cryptobranchus alleganiensis*) mainly produce cortisol in response to a stressor (Hopkins et al., 2020). Yet, the vast majority of studies on a popular aquatic model species, the clawed frog (*Xenopus* sp.), measure corticosterone (e.g., Jaudet and Hatey, 1984; Kulkarni and Buchholz, 2012; Shewade et al., 2020), likely based on foundational *in vitro* studies with *Xenopus laevis* interrenal cells (Chan and Edwards, 1970; Jolivet-Jaudet and Leloup-Hatey, 1984). Conversely, in samples from dermal swabs, cortisol increases in response to ACTH in terrestrial green treefrogs (*Hyla cinerea*), American toads (*Anaxyrus americanus*), and red-spotted newts (*Notophthalamus viridescens*) (Santymire et al., 2018). Further equivocal results about cortisol and corticosterone in amphibians, including contradictory results in the same species (such as *Xenopus laevis* and *Rana pipens*), are nicely summarized in Hopkins et al. (2020).

In sum, contradictory findings call into question whether corticosterone can be assumed to be the only GC with biological importance in amphibians, especially as we expand our study of endocrinology to non-traditional species. Measuring both GCs will help us discern the role of different GCs across development and associated with distinct life histories, and will build a more complete picture of amphibian physiology and health in captivity and in the wild.

### 1.3 Stress physiology in Neotropical poison frogs

Neotropical poison frog (Dendrobatidae) species have long garnered interest from evolutionary biologists and hobbyists, and have gained recent attention in behavioral ecology and neuroethology due to their interesting parental behavior and spatial cognition (Burmeister, 2022; Pašukonis et al., 2014; Peignier et al., 2024; Ringler et al., 2023; Roland and O’Connell, 2015; Westrick et al., 2023). These often brightly colored, primarily diurnal, terrestrial frogs are threatened by agrochemicals (Angulo et al., 2019), climate change (Whitfield et al., 2007), smuggling (Auliya et al., 2016), habitat destruction or fragmentation (Becker et al., 2007; Cushman, 2006), and susceptibility to the amphibian chytrid fungus *Batrachochytrium dendrobatidis* (*Bd*) (Courtois et al., 2015). Due to their conservation status (Guillory et al., 2019) the development of non-invasive measurements of GCs is important for studying poison frog physiology and informing conservation efforts. Because poison frogs are generally small, with an average snout-vent length (SVL) of ∼17-60 mm, repeated plasma sampling is not feasible, and there is a need to further develop non-invasive methods of studying the endocrinology of poison frogs in the wild and the lab.

Our goal for this study was to characterize GC abundance and ACTH-responsivity in adult dyeing poison frogs, *Dendrobates tinctorius*. *D. tinctorius* has received growing interest in behavioral and evolutionary studies in recent years (Mayer et al., 2025; Moskowitz et al., 2022; Pašukonis et al., 2022; Rojas and Pašukonis, 2019; Sabino-Pinto et al., 2024; Schlippe Justicia et al., 2024; Soto et al., 2024). In the context of hormones and behavior, we previously found corticosterone is more abundant and ACTH-responsive in *D. tinctorius* tadpoles (Surber-Cunningham et al., 2024), but also found a surprising role for cortisol in parental behavior (Fischer and O’Connell, 2020).

To build a more complete picture of poison frog physiology, we used non-invasive waterborne hormone sampling to allow for repeated measures from the same individuals (e.g. Baugh et al., 2018; Gabor et al., 2013; Rodríguez et al., 2022) and an ACTH challenge to assess the responsivity of the HPA axis to ACTH. We then compared glucocorticoid concentrations between water and plasma samples from the same individuals to assess whether water samples are representative of circulating hormone levels in blood. To test if the patterns we found were consistent across Dendrobatids, we also characterized cortisol and corticosterone excretion in four additional, related poison frog species. All together this study shows how the quantification of multiple GCs can uncover potentially important variation that is more complex than just the ‘stress’ physiology of this particularly vulnerable clade of frogs and contributes pieces to the puzzle of GC physiology in amphibians and vertebrates more broadly.

## 2. Materials and methods

### 2.1 Animal husbandry

We housed all frogs in our breeding colony at University of Illinois Urbana-Champaign in glass terraria (18x18x18”, 12x12x18”, or 36x18x18” Exo Terra®) with soil, sphagnum moss, live tropical plants, and coconut shelters. We kept frogs on a 12:12 h light cycle and fed them flightless *Drosophila* fruit flies (*D. melanogaster* or *D. hydei*) dusted with vitamin supplements three times weekly. Our automated RO water misting system kept tank humidity >75% and the room was kept at 21-24°C. In addition to *D. tinctorius*, we measured GCs in golden poison frogs (*Phyllobates terribilis*), mimic poison frogs (*Ranitomeya imitator*), Zimmermann’s poison frogs (*Ranitomeya variabilis*), and Anthony’s poison frogs (*Epipedobates anthonyi*). These species share many behaviors, including parental care, and are endemic to tropical regions of Central and South America, but vary in body size and toxicity in the wild (Wells, 2007). *P. terribilis* and *D. tinctorius* are among the largest poison frogs (∼37-60 mm snout-vent length (SVL)), *E. anthonyi* are intermediate (∼19 to 25 mm SVL), and both *Ranitomeya* are among the smallest (∼17-22 mm SVL). We housed adult frogs in groups of two to three individuals of the same species, either one or two males with one female (*E. anthonyi*, *D. tinctorius*, *R. variabilis*, *R. imitator*) or as a group of two to three frogs that were too young to be sexed at the time of sampling (*P. terribilis*). All frogs were at least 1 year old. For dosing during the ACTH challenges in *D. tinctorius*, we weighed frogs prior to sampling by placing them in a plastic container on a scale and subtracting the weight of the container. All procedures were approved by UIUC Institutional Animal Care and Use Committee (protocol #20147).

### 2.2 Hormone sampling

To collect waterborne hormones, we placed individual frogs in glass containers with room temperature RO water based on size. For the smaller poison frogs (*E. anthonyi*, *R, variabilis*, and *R. imitator*), we used 4 cm x 4 cm x 5 cm hexagonal glass jars with lids filled with 15 mL of RO water. For the larger frogs (*D. tinctorius* and *P. terribilis*), we used 6.5 cm x 6.5 cm x 6 cm glass tubs with lids and 40 mL of RO water. Our goal was to use enough water to cover frogs’ highly vascularized pelvic skin in a container small enough that the frogs would stay in the water and not climb the walls. We left frogs in the water for one hour before returning them to their home terraria or a new water tub for additional sampling. We collected all samples 4.5-8 hours into the frog’s 12 h light cycle. We filtered water samples with Whatman® 1 filter papers to remove any large particulates, prior to steroid hormone extraction (see Section 2.4). To compare across species, we collected water samples from *D. tinctorius*, *E. anthonyi*, *P. terribilis*, *R. imitator*, and *R. variabilis*). We stored all water samples at −20°C until extraction. For the plasma and waterborne hormone comparison, we collected water samples as described above for *D. tinctorius* immediately prior to collecting plasma. We subsampled half of the water from n = 155 water samples for additional projects and stored each subsample separately. To collect plasma samples, we anesthetized frogs with benzocaine (Orajel™) and euthanized via rapid decapitation. This process took <5 minutes, in line with generally accepted time courses for plasma sample collection. We used heparinized microhematocrit capillary tubes to collect trunk blood from the body. We combined all the collected blood from an individual in a heparinized microcentrifuge tube and centrifuged for 5 min at 5,000 rpm to separate the plasma. Using a micropipette, we collected the plasma supernatant and stored all plasma samples at −20°C.

### 2.3 ACTH hormone challenge

For the ACTH hormone challenges, we used a within-individual repeated measures design with 18 adult *D. tinctorius* (9 male, 9 female) for a total of 36 challenges (18 ACTH injections and 18 control injections). We injected frogs with ACTH (Sigma-Aldrich®, product no. A7075) diluted in 0.9% saline and a vehicle injection of only 0.9% saline with 6 days between injections. The order of injections was counterbalanced across individuals such that half of the frogs were first injected with vehicle, and half were first injected with ACTH. Prior to the start of trials, we dissolved the ACTH in saline at a concentration of 0.125 ug/µL and aliquoted the solution into insulin syringes which were frozen at −20°C until the day of injection, along with saline vehicle aliquots. On the day of the trial, we thawed the aliquots of saline and ACTH and calculated the volume needed to inject each frog with a dosage of 0.5 ug ACTH per g body weight (mean ± SD = 5.89 ± 1.5 g), based on previous work using ACTH challenges in Neotropical frogs (Baugh et al., 2018). Since the precision of the insulin syringes was limited, we rounded to the nearest 10 µL to get the volume for injection with both ACTH and saline. For each trial, we collected a waterborne hormone sample, as described above, for one hour prior to injection as our “pre-injection” sample. We then used insulin syringes to inject frogs intraperitoneally. After injecting, we immediately placed the frogs in a new water bath for one hour to collect the “post-injection” sample. After the post-injection sample, we returned frogs to their home terraria. We repeated this process the following week with the alternate treatment (ACTH or saline). We performed all trials between five and seven hours after the lights turned on. We split each water sample (four samples per 18 frogs = 72 samples total) in half for additional projects and stored subsamples separately.

### 2.4 Steroid hormone extraction

For all water samples, we used C18 cartridges (Waters™ Corporation, Milford, MA) for solid-phase extraction to collect total steroid hormones. These cartridges are widely used for water hormone extraction for fish and frogs (Baugh et al., 2018; Gabor et al., 2013; Gabor and Grober, 2010; Kim et al., 2018). To test the recovery rate of the cartridges, we spiked clean water samples with a known quantity of hormone and found a 101% and 91% average recovery of cortisol and corticosterone, respectively. With cartridges on a vacuum manifold, we used a vacuum pump to prime the filters with 6 mL 100% methanol. We then rinsed the cartridges with 6 mL RO water before passing the sample through the cartridge. We eluted the hormones from the cartridges with 4 mL of 100% methanol into 13x100mm borosilicate glass vials. We dried samples under a stream of nitrogen gas in a 37°C dry heating block until completely dry. We stored desiccated samples at −20°C until reconstituted with assay buffer for enzyme linked immunosorbent assay (ELISA) measurement of total steroid hormones (see Section 2.6).

To measure glucocorticoids from plasma samples, we used the dissociation reagent included in our ELISA kits (see Section 2.6). Due to the limited volume of plasma collected from each frog, we chose to use the dissociation reagent over doing an extraction process due to small sample volumes and to avoid losing sample through the extraction steps. We combined 5 µL of dissociation reagent and 5 µL of plasma, vortexed, and incubated for a minimum of 5 min at room temperature. We then added 450 µL of assay buffer to make a 1:92 dilution before running samples on ELISA plates.

### 2.6 Enzyme linked immunosorbent assays

We used DetectX® ELISA kits from Arbor Assays™ (Ann Arbor, MI) to measure total cortisol and corticosterone concentration in plasma and water samples. We used each kit according to the manufacturer’s instructions except for using the same assay buffer (catalog no. X065) for both kits to allow us to split samples across cortisol and corticosterone plates, following manufacturer recommendations. For the corticosterone kit, we followed instructions for the 100 µL format because the standard curve for the 100 µL format can measure smaller concentrations than the 50 µL format. For both cortisol and corticosterone, we measured the optical density at 450 nm. After removing samples with a coefficient of variation (CV) > 20% (n = 77 corticosterone and n = 62 cortisol, out of 241 measurements each), the average intra-assay CV across the remaining samples for the analyses was 8.2% for cortisol and 8.1% for corticosterone. The cross reactivity for the cortisol ELISA plate with corticosterone is reported by the manufacturer as 1.2% and for the corticosterone ELISA plate cross reactivity with cortisol is reported as 0.38%. Information about cross reactivity with other molecules can be found on the Arbor Assays™ website.

We used a four-parameter logistic curve (4PLC) regression to fit the respective standard curve on each plate using MyAssay software recommended by the plate manufacturer (www.MyAssay.com). Based on this curve, we calculated the concentration of cortisol and corticosterone (pg/mL of assay buffer) for each sample and averaged across duplicates. Due to low concentrations of both GCs in some species, some samples were too low to fall on the standard curve (n = 13 corticosterone and n = 30 cortisol, out of 241 measurements each), though some had concentrations of cortisol that were too high (n=10). To be conservative, for samples that were off the standard curve in either direction (too low or too high), we censored the data at the minimum or maximum detection limits of the assay, respectively. To confirm the censored data did not skew our findings, we ran additional analyses with only samples measuring within the assay detection limits and CV < 20%, and the overall pattern of results remained the same.

We report release rate (pg/hr) for both cortisol and corticosterone from water samples (Gabor et al., 2013). For samples that were split in half prior to extraction, we multiplied the rate (pg/hr) by two to allow comparison with samples that were not split.

We tested each assay for parallelism by combining 24 additional water samples from adult *D. tinctorius* to create a highly concentrated pooled sample and then serial diluting by 1/2 of the reconstituted extracts for a dilution curve with seven samples in total. This serial dilution sample was split four ways to test parallelism across four different ELISA plates for steroid hormones, including cortisol and corticosterone. We confirmed parallelism by comparing the slope of the serially diluted curve to the slope of the standard curve using an interaction term in a linear model. We found the dilution curves were not significantly different from the standard curve for their respective hormone (cortisol: ß=-0.34, SE = 1.08, p = 0.76; corticosterone: ß=-5.09, SE = 3.06, p = 0.12).

### 2.7 Statistical analysis

We used R v4.2.2 (R Core Team, 2023) in RStudio v3.1.446 (Posit Team, 2023) for all statistical analyses and ran all analyses on log transformed hormone release rates to normalize residuals.

First, we compared the concentration of cortisol and corticosterone in plasma and water from *D. tinctorius*. We calculated the original plasma sample concentration by multiplying the concentration measured in the assay by the dilution factor. For the analyses with water samples alone, we included all unmanipulated samples, including those taken immediately prior to euthanasia and samples taken on the first day of trials before the injection during ACTH challenges. We fit two separate linear models (R packages lme4 (Bates et al., 2015) and lmerTest (Kuznetsova et al., 2016)) with concentration (pg/mL) in plasma (n = 32 samples, 17 frogs) and hormone release rate (pg/hr) in water samples (n = 59 samples, 32 frogs). For both models, we included GC type (cortisol or corticosterone) and sex as fixed effects and frog ID as a random effect to control for repeated samples, since both hormones were measured in each sample.

Next, we were interested in whether the GC types are correlated with one another which would suggest measuring either one is sufficient in most scenarios. To assess the correlation between GC types, we fit two linear models (plasma model, n = 15 samples; water model, n = 23 samples) with cortisol concentration or release rate as the response variable and corticosterone as the fixed effect. Additionally, to examine the relationship between waterborne glucocorticoids and circulating glucocorticoids in plasma, we fit linear models for both cortisol (n = 11 frogs) and corticosterone (n = 16 frogs), separately, with plasma concentrations (log transformed pg/mL) as the fixed effect predicting water hormone release rate (log transformed pg/hr) for the matched plasma-water samples.

For the ACTH challenges, we fit a linear mixed effects model to ask if there was an effect of treatment (ACTH or control) on either cortisol or corticosterone. The model had the following main effects: SVL in cm, GC type (cortisol or corticosterone), time point (pre- or post-injection), and injection type (ACTH or saline). Because these data were all collected from one species, we included SVL in the model to account for differences in hormone release rate due to body size within species, as is recommended in waterborne hormone sampling in fish (Scott and Ellis, 2007). We chose SVL over mass to avoid variation in hydration levels which can have a substantial effect on body mass in amphibians. We also included a three-way interaction between GC type, time point, and injection type. This interaction indicated whether the difference between pre- and post-injection varied by injection type and if this relationship varied by GC type. In other words, was there a change in cortisol from pre- to post-injection based on injection type and was this change (or lack thereof) the same or different in corticosterone? We included trial ID as a nested random effect within frog ID to account for repeated measures across trials and individuals. We used post-hoc pairwise comparisons using the multcomp package (Hothorn et al., 2008) to interpret interaction effects. After filtering for CV, our sample size was 26 pre- and 26 post-injection cortisol measurements and 27 pre- and 30 post-injection corticosterone measurements. Because our main interest was how the within-individual response to the injection varied across injection types, all four samples for each individual were run on the same plate to reduce possible confounding effects of inter-plate variation within individuals and trials.

Across species, we were primarily interested in whether the release rates of the two GCs differed within species, rather than absolute GC differences between species. To this end, we fit a linear mixed model for log transformed hormone release rate with a fixed effect of the interaction between species and glucocorticoid type (corticosterone vs cortisol) with sample ID as a random effect. We ran a Type III ANOVA on the linear mixed model to assess the overall effect of the interaction term. We then ran post-hoc simultaneous pairwise comparisons using the multcomp package (Hothorn et al., 2008) to specifically look at the difference in release rate between cortisol and corticosterone within species (e.g., *P. terribilis* cortisol vs. *P. terribilis* corticosterone). We did not include SVL as a correction for body size, as large species differences in average body size confound the effect of SVL and species if both are included in the same model. Furthermore, SVL was not significantly correlated with GCs in our *D. tinctorius* specific analyses (Table 1), which is consistent with previous studies on waterborne hormones in frogs (e.g. Gabor et al., 2013). We also did not include sex as a covariate since not all individuals could be confidently sexed at time of sampling. However, in a reduced model with only individuals that we could confidently sex, we found no overall effect of sex on GC production (F_1,79.5_ = 0.033, p = 0.86). We were primarily interested in comparing the abundance of GCs within species rather than across species. Our final species comparison dataset included 59 measurements from *D. tinctorius* (32 corticosterone, 27 cortisol), 37 measurements from *E. anthonyi* (15 corticosterone, 22 cortisol), 23 measurements from *P. terribilis* (8 corticosterone, 15 cortisol), 35 measurements from *R. imitator* (16 corticosterone, 19 cortisol), and 19 measurements from *R. variabilis* (3 corticosterone, 16 cortisol). While we had few measurements of corticosterone in *R. variabilis* samples, we include them in the analysis as valuable preliminary information about a species that, to our knowledge, has no current published glucocorticoid data.

**Table 1.** Results from linear model for hormone release rate across injection type and time points. The reference groups are corticosterone, control (saline) injection, and pre-injection time point. P values < 0.05 are listed in bold.

| log(release rate, pg/hr) |  |  |  |
| --- | --- | --- | --- |
| Predictors | Estimates | CI | p |
| intercept | 4.93 | 3.54 – 6.32 | **<0.001** |
| snout-vent length (SVL, cm) | 0.09 | -0.22 – 0.39 | 0.581 |
| glucocorticoid type (cortisol) | -0.44 | -0.87 – -0.01 | **0.044** |
| injection type (ACTH) | 0.40 | -0.09 – 0.88 | 0.105 |
| time point (post) | -0.59 | -1.02 – -0.17 | **0.007** |
| cortisol x ACTH | 0.06 | -0.54 – 0.66 | 0.834 |
| cortisol x post-injection | 0.62 | 0.01 – 1.23 | **0.047** |
| ACTH x post-injection | 0.96 | 0.38 – 1.55 | **0.001** |
| cortisol x ACTH x post-injection | -0.98 | -1.81 – -0.14 | **0.023** |
| **Random Effects** |  |  |  |
| $\sigma^2$ | 0.29 | | |
| T00 trial:frogID | 0.12 |  |  |
| T00 frogID | 0.02 |  |  |
| ICC | 0.33 |  |  |
| N<sub>trial</sub> | 35 |  |  |
| N<sub>frogID</sub> | 18 |  |  |
| Observations | 109 |  |  |
| Marginal R<sup>2</sup> / Conditional R<sup>2</sup> | 0.314 / 0.541 |  |  |

Finally, we compared the relationship between cortisol and corticosterone across species. Due to our limited sample sizes for the smaller *Ranitomeya*, we had too few samples with data for both cortisol and corticosterone release rate, so we restricted the comparison to the larger species (*D. tinctorius*, *P. terribilis*, and *E. anthonyi*). To compare with the *D. tinctorius* model (above), we fit separate linear models for *P. terribilis* (n = 7) and *E. anthonyi* (n = 12) with cortisol release rate as the response variable and corticosterone as a fixed effect.

## 3. Results

### 3.1 Glucocorticoid abundance and correlations in D. tinctorius

In *D. tinctorius*, corticosterone was statistically more abundant than cortisol in both plasma (ß= −2.04, p < 0.001; Fig 1B) and water samples (ß= −0.55, p < 0.001; Fig 1C). In plasma samples, corticosterone concentration was 22.8% higher than cortisol on average. In water samples, the difference between GC types was smaller; corticosterone release rate was 11.4% higher than cortisol on average.

When we compared water with plasma samples in *D. tinctorius*, the concentration of cortisol in plasma was positively correlated with the release rate of cortisol in water (ß=1.68, p = 0.037; Fig 1D). We found a slightly positive but statistically non-significant relationship between corticosterone concentration in plasma and corticosterone release rate in water (ß=0.27, p = 0.39; Fig 1D).

In both plasma and water, cortisol and corticosterone were positively correlated, but this relationship was weaker in plasma samples and only statistically significant in water samples (plasma: ß= 0.17, p = 0.15; water: ß= 0.45, p = 0.0077; Fig 2).

**Figure 2.**
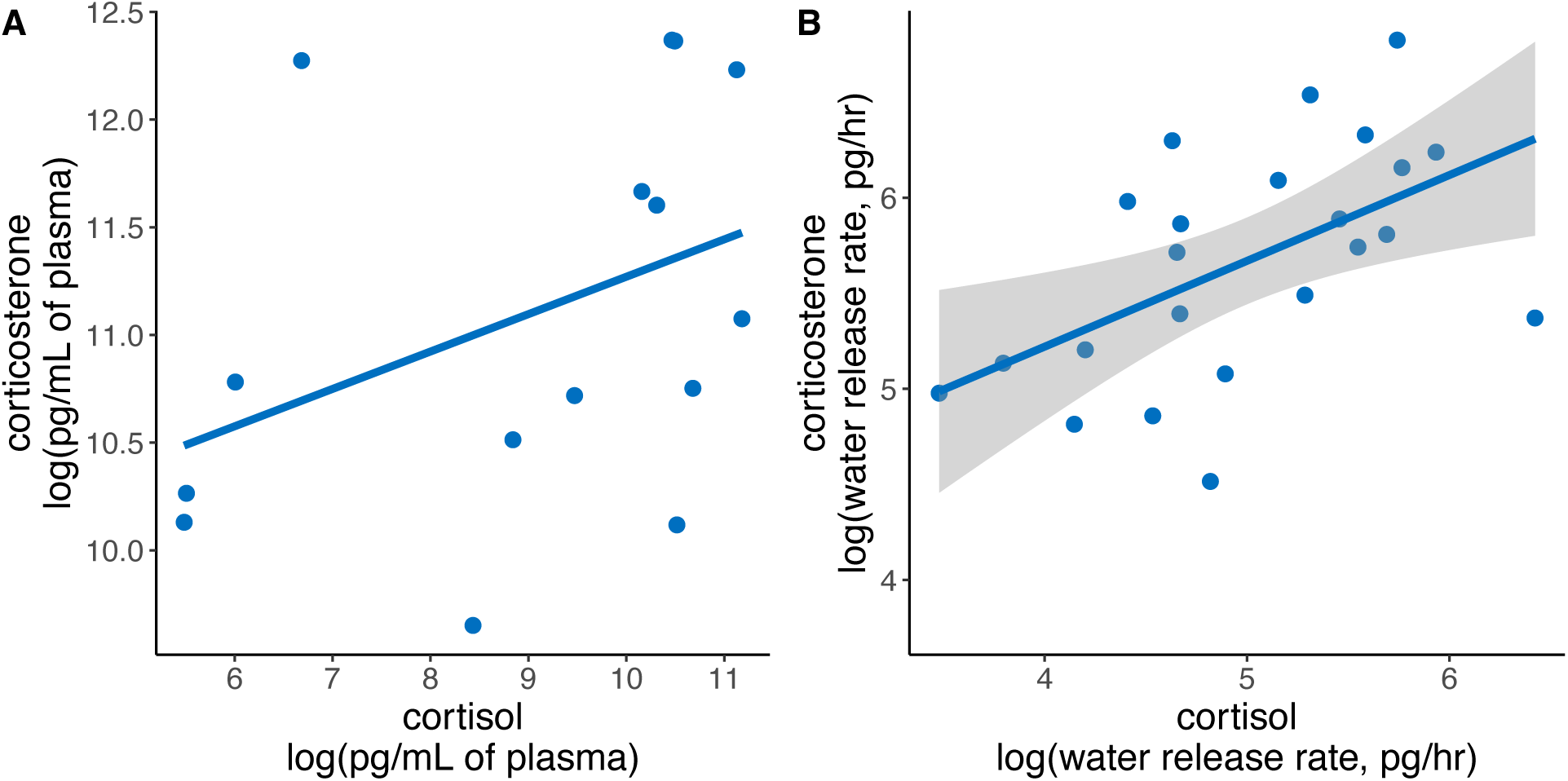
Correlation between glucocorticoids. Corticosterone and cortisol were positively correlated in both (A) plasma and (B) water. However, the relationship was only statistically significant in (B) water samples. Grey shaded area is the confidence interval around the statistically significant linear regression for water samples.

### 3.2 ACTH challenges

The effect of an injection (ACTH vs vehicle) varied by GC and injection type (GC x injection type x time point: ß= −0.97, p = 0.024; Table 1; Fig 3). We found overall decreased GC release rates post-injection (ß=-0.59, p = 0.007; Table 1) driven by a decrease in corticosterone in saline injected frogs (pairwise comparison of corticosterone between pre- and post-injection saline: ß = 0.59, z = 0.21, p = 0.031; Fig 3). After an injection, frogs had higher corticosterone release rates when injected with ACTH compared to saline alone (pairwise comparison of corticosterone between ACTH and saline post-injection: ß = 1.36, z = 5.86, p <0.001; Fig 2).

**Figure 3.**
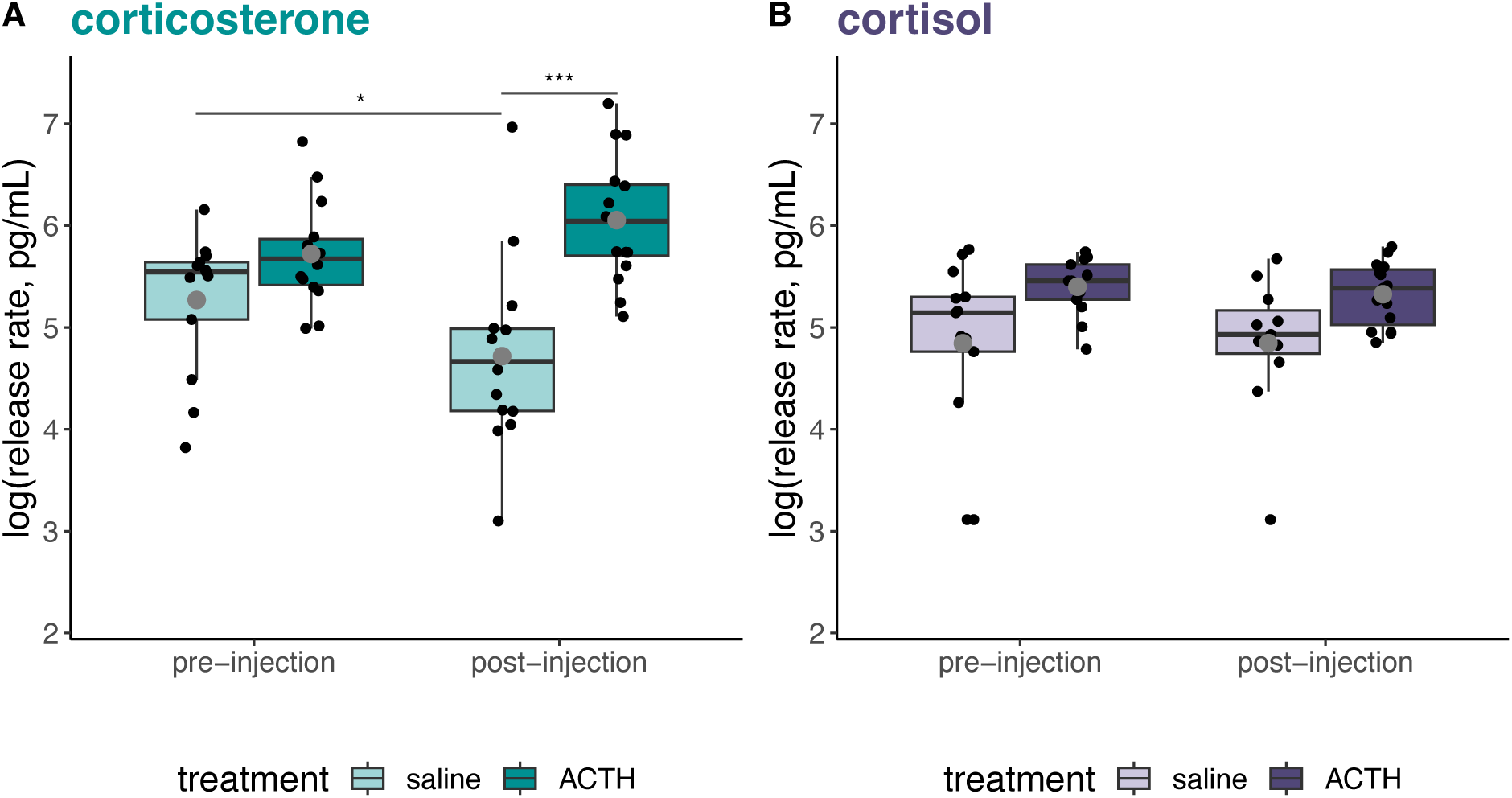
Glucocorticoid response to an ACTH injection compared to a saline control injection. (A) Corticosterone release rate was higher after an ACTH injection than after a saline injection. (B) In contrast, we found no difference in cortisol release rate between samples taken before and after injection of either saline or ACTH. Purple boxplots indicate cortisol data. Boxplots for corticosterone data are in teal. Lighter colors (on the left within injection groups) indicate the control/saline injection for the respective glucocorticoid. Grey dots indicate the mean for each treatment group. Black dots are individual water samples. (* = p < 0.05, *** = p < 0.001)

Cortisol release did not vary pre- and post-injection for either type of injection (pairwise comparisons of cortisol between pre- and post-injection: saline ß = −0.02, z = −0.09, p = 1.00, ACTH ß = −0.03, z = −0.03, p = 1.00; Fig 3).

### 3.3 Species variation in relative glucocorticoid abundance

We found species differed in their release rate of each glucocorticoid (F_4,71.8_=9.57, p < 0.0001; Fig 4). Specifically, pairwise comparisons within species revealed *D. tinctorius* released slightly more corticosterone (ß = −0.55, p = 0.053), while three species (E*. anthonyi*, *R. imitator*, and *R. variabilis*) released more cortisol (Table 2; Fig 4). In *P. terribilis*, there was a trend for higher cortisol release than corticosterone, but the difference was not statistically significant (Table 2; Fig 4). In a less conservative analyses, excluding all samples that did not fall within the range of the assays (i.e., remove censored values), the overall patterns were the same, however the difference between GC types became statistically significant for *P. terribilis* (ß = 0.98, p = 0.020), which released 2.5x more cortisol than corticosterone.

**Figure 4.**
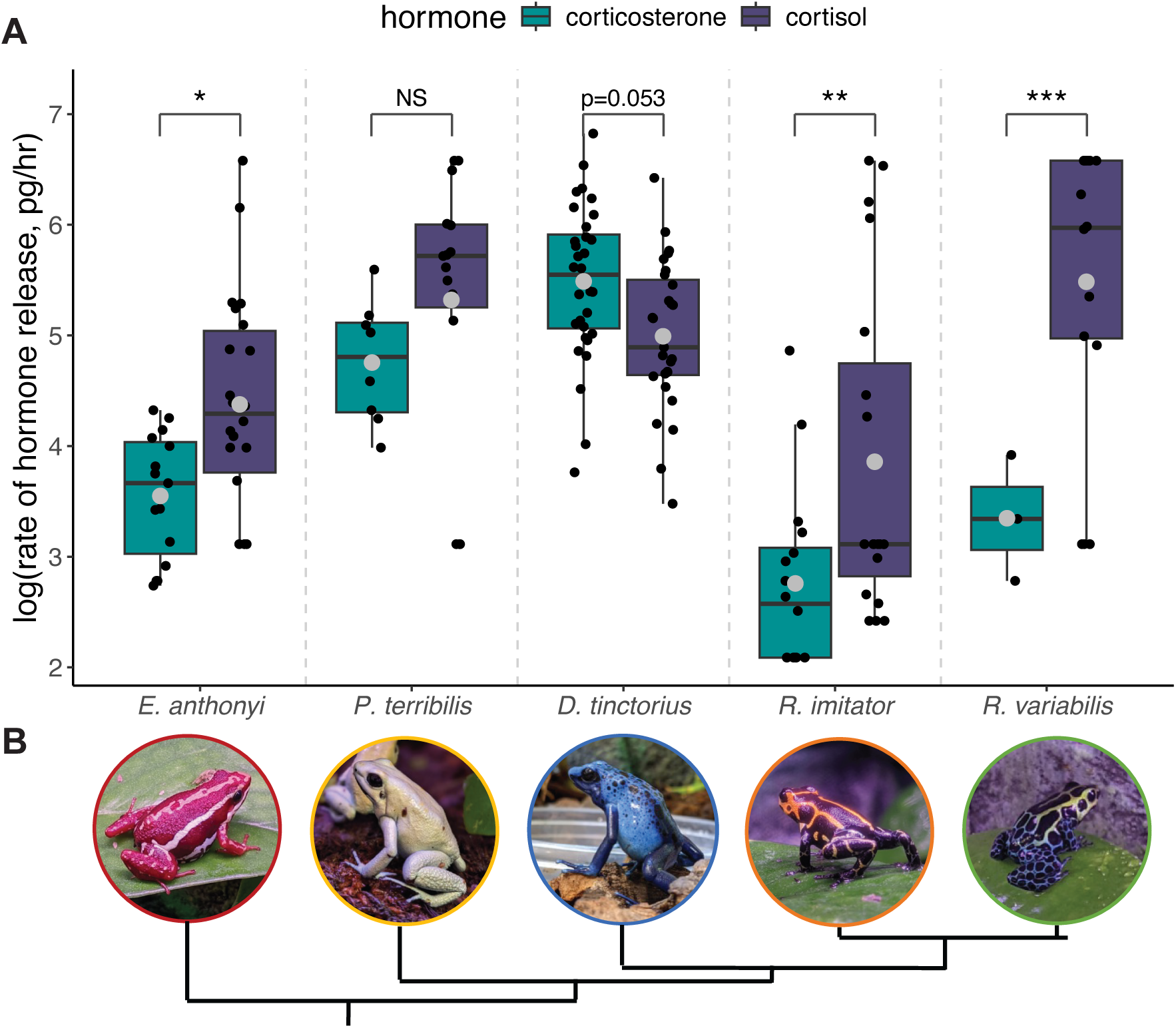
Species variation in relative glucocorticoid abundance. (A) For four out of the five species tested, the release rate of corticosterone was lower than the release rate of cortisol. In *D. tinctorius*, corticosterone was released 1.64 times more than cortisol. Boxplots for corticosterone are teal (left) and for cortisol are purple (right). Grey dots indicate the mean for each group. Black dots represent individual samples (NS = not significant, * = p < 0.05, ** = p < 0.001, *** = p < 0.0001). (B) The species names correspond to the respective photos beneath the x-axis labels. The tree below the photos indicates the phylogenetic relationships between species (Guillory et al., 2019).

**Table 2.**
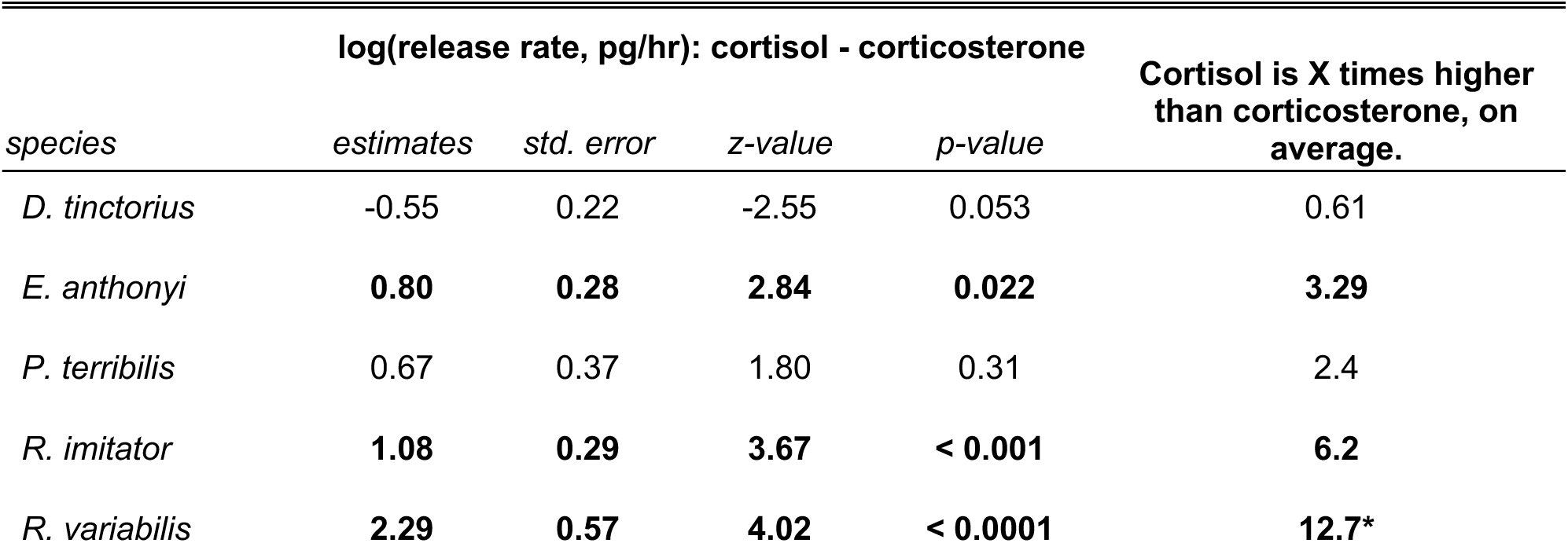
Results from simultaneous multiple pairwise comparisons of means for hormone release rate across species. P-values < 0.05 are in bold. *The difference for *R. variabilis* should be interpreted with caution due to small sample sizes.

Unlike the positive correlation between cortisol and corticosterone in water samples from *D. tinctorius* (see Section 3.1), the two GCs were not statistically significantly correlated in *P. terribilis* (ß = 0.24, p = 0.53; Fig 5) and *E. anthonyi* (ß = 0.29, p = 0.65; Fig 5).

**Figure 5.**
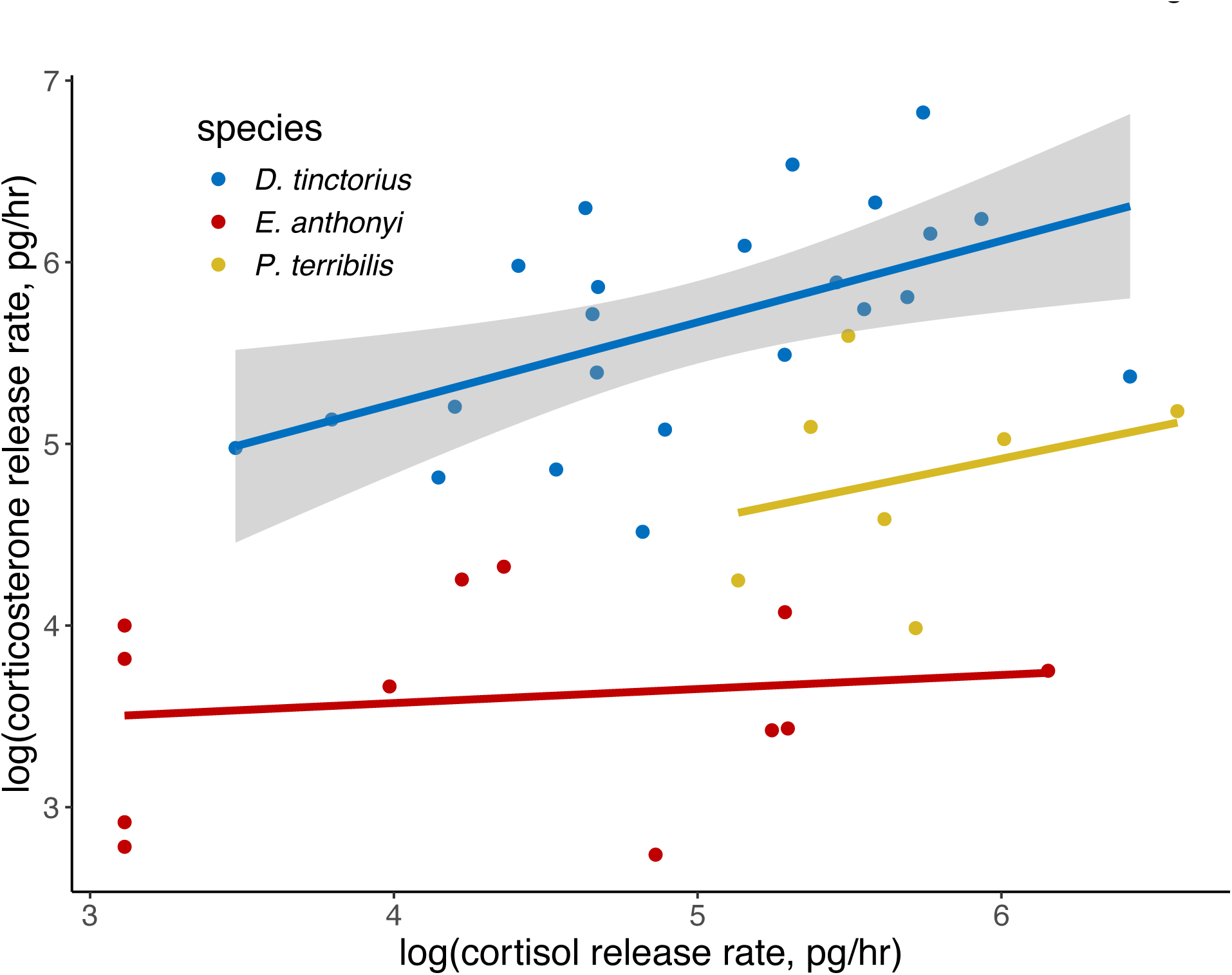
Species variation in the relationship between GC types. The positive relationship between cortisol and corticosterone was stronger in water samples. Grey shaded area is the confidence interval around the statistically significant linear regression for *D. tinctorius* water samples. *D. tinctorius* data are the same as those in Fig 2. We added them here as well for ease of comparison across the species.

## 4. Discussion

Glucocorticoids are involved in many physiological, developmental, and behavioral processes that are essential for survival and fitness (MacDougall-Shackleton et al., 2019; Nicolaides et al., 2015); therefore, understanding glucocorticoids and HPA axis dynamics can inform research and conservation efforts and improve our understanding of how animals are impacted by and respond to environmental changes and threats (Busch and Hayward, 2009; Millspaugh and Washburn, 2004). Our initial goal was to identify which glucocorticoid was most abundant and responsive to activation of adrenocortical tissue via injection of ACTH, i.e., the dominant GC by the typical definition, in our primary focal species, *D. tinctorius*. Our interest in the question was spurred generally by an increasing awareness of the subtleties of hormonal effects within and across species, and specifically by our previous findings that increased cortisol is associated with parental care in adult *D. tinctorius* (Fischer and O’Connell, 2020), yet corticosterone is more abundant and ACTH-responsive in *D. tinctorius* tadpoles (Surber-Cunningham et al., 2024). Given both GCs were measurable in *D. tinctorius*, we further confirmed non-canonical patterns in adults of four additional Dendrobatid species. Our findings challenge the common assumption that corticosterone is always more abundant in frogs and suggest that focusing only on the ACTH-responsive GC can overlook potentially important ACTH-independent regulation of other physiologically important GCs. Below we discuss the biological implications, remaining technical challenges and limitations, and prospects of this work.

### 4.1 Glucocorticoid abundance and dominance in D. tinctorius

We used non-invasive waterborne hormone sampling to quantify cortisol and corticosterone excretion rates in adult *D. tinctorius* poison frogs. In many other species, including frogs and *D. tinctorius* tadpoles (Surber-Cunningham et al., 2024), one GC is substantially more abundant.

For example, cortisol concentration is around 10x higher than corticosterone in eastern hellbender plasma (Hopkins et al., 2020), likewise corticosterone is around 10x higher than cortisol in rat serum (Gong et al., 2015). In comparison, in *D. tinctorius* water samples, corticosterone was less than 2 times as abundant as cortisol. As a substantially greater abundance often predicts ACTH responsivity, this small difference between cortisol and corticosterone and previous evidence for an association between cortisol and behavior in adult *D. tinctorius* (Fischer and O’Connell, 2020) left open the question of GC dominance by the typical definition of ACTH responsivity.

In response to an ACTH challenge, we found evidence for corticosterone as the “dominant” GC in *D. tinctorius*. The release rate of corticosterone decreased following a saline injection, regardless of whether the saline injection was the first or second injection the frogs experienced. This decrease suggests frogs were no longer mounting a stress response due to the sampling process; however, an ACTH injection prevented this decrease in corticosterone, demonstrating a response relative to controls (Fig 3). Interestingly, cortisol did not respond to either an injection of ACTH or saline (Fig 3) suggesting cortisol is not responsive to acute adrenal activation and does not fluctuate across short time periods (i.e., does not show an increased or habituation to waterborne sample collection). In combination, these results indicate corticosterone is more responsive to acute adrenal activation than cortisol, despite similar amounts of cortisol and corticosterone prior to injections. Often, the GC that responds most to adrenal activation through an ACTH challenge is also more abundant (e.g., Forsburg et al., 2019; Hopkins et al., 2020).

While this pattern of abundance relating to ACTH responsiveness holds in *D. tinctorius*, the difference in abundance between the GCs was relatively small compared to other species. For example, across 18 species of mammals cortisol is more responsive to ACTH and cortisol concentrations were between 7.5 and 49 times higher than corticosterone (Koren et al., 2012). Similarly, in the American bullfrog (*R. catesbeiana*), corticosterone is ACTH responsive (Krug et al., 1983) and ∼26 times as abundant than cortisol (Krug et al., 1983; Wright et al., 2003). Our finding that corticosterone and cortisol are similar in abundance yet differentially responsive to ACTH underscore the need to characterize both GCs and raise questions as to their primary roles in poison frog physiology and behavior.

Evidence for corticosterone as the ACTH-responsive GC in *D. tinctorius* aligns with our findings in *D. tinctorius* tadpoles (Surber-Cunningham et al., 2024), a previous study using restraint stress in the mimic poison frog (*R. imitator*; Nowicki et al., 2024), and broader taxonomic trends in amphibians. However, our previous findings also suggest a functional role for cortisol in parental care (Fischer and O’Connell, 2020). We hypothesize that corticosterone and cortisol may serve distinct functions and may be released via distinct mechanisms depending on context, thus providing a means for modular regulation of behavioral and physiological functions of hormones (McGlothlin and Ketterson, 2008). Though speculative, this hypothesis aligns with evidence for distinct roles of the two GCs in different tissues (Schmidt and Soma, 2008) and due to differential binding affinities with their shared receptors (Katsu et al., 2016; Sutanto and De Kloet, 1987). There are various non-ACTH routes of GC production, including regulation via neurotransmitters and neuropeptides involved in behavior (Bornstein and Chrousos, 1999), which provide a potential mechanism for modularity in GC release. Further, while both GCs bind to the same receptor types, the potential for differential release mechanisms poses an intriguing reframing of our understanding of how the HPA axis functions. For example, the parental state involves a myriad of physiological changes (Numan, 2020; Rosenblatt and Snowdon, 1996) and perhaps the long-term physiological versus short-term behavioral demands of parenting rely on the production and localization of glucocorticoids through different routes. We suggest future studies explore the idea that ACTH and ACTH-independent mechanisms could provide modular regulation of responses to various stressors and stressful contexts, such as parental care.

Though our interpretations remain speculative, there is growing precedent for the differential functionality of cortisol and corticosterone within species (Schmidt and Soma, 2008; Vera et al., 2017). In some species and contexts, cortisol is more responsive to acute stress than corticosterone (Koren et al., 2012; Venkataseshu and Estergreen, 1970; Vera et al., 2012, 2011), however the two GCs may also simultaneously elevate in more chronic stress states with high energetic burdens. For instance, both cortisol and corticosterone increase during pregnancy in humans (Wintour et al., 1978), little brown bats (*Myotis lucifugus,* Reeder et al., 2004), North Atlantic right whales (Hunt et al., 2017), and Asian elephants (*Elephas maximus*, Kajaysri and Nokkaew, 2014). In essence, the ratio of the two GCs in circulation may be dependent on physiological and/or behavioral state. In the context of increased cortisol during parenting in *D. tinctorius* (Fischer and O’Connell, 2020), an increase in corticosterone, but not cortisol, following an ACTH injection suggests non-ACTH routes of GC production may be important in upregulating cortisol during behavioral states with higher energetic demand, such as tadpole transport. Such modular regulation could occur through changes in the regulation of the enzyme required to produce cortisol (17α-hydroxylase). Further studies are needed to explore these intriguing possibilities.

The motivation to measure both glucocorticoids is further underscored by our results, and the results of others (e.g. Koren et al., 2012), which suggest that the two GCs are not always correlated with one another nor entirely redundant in function. In non-invasive water samples from *D. tinctorius*, cortisol was positively correlated with corticosterone. However, this correlation was not significant in plasma samples or in water samples from *P. terribilis* or *E. anthonyi*. These results suggesting the ratio of circulating GCs may fluctuate, as discussed above, and the relationship between the GCs may vary by sampling matrix. One possibility yet to be explored is the temporal variation in the regulation of each GC. If the abundance of one GC is more dynamic over short time periods than the other, this could lead to a more stable positive relationship between the GCs in cumulative samples, such as water, than the more immediate sampling of plasma.

### 4.2 Species variation in glucocorticoids

Following our unexpected results in *D. tinctorius*, we characterized waterborne hormone levels in four additional poison frog species, yielding data from five species representing four genera. Results from all five species across the *Dendrobatidae* family contradict the common assumption that corticosterone is the more abundant GC in amphibians (Moore and Jessop, 2003; Norris and Carr, 2013; Wingfield and Romero, 2011), and in particular in terrestrial amphibians (Jungreis et al., 1970). In waterborne hormone samples, we found cortisol was released at a higher rate than corticosterone in all four additional species, though magnitude varied (2.4-12.7x; Fig 4). We opportunistically chose these five species of poison frogs representing four genera.

More systematic sampling is needed to determine whether variance in relative GC abundance across Dendrobatids is associated with phylogeny, life history, or some combination. Nonetheless, we can confidently say that this group does not follow the general assumption of higher corticosterone abundance in (terrestrial) amphibians. Most studies comparing cortisol and corticosterone within-species have been done in mammals, with few systematic studies in other taxa (Vera et al., 2017). The few studies in amphibians that have looked at both cortisol and corticosterone report a range of empirical results across different sample types with no consistent pattern, as discussed above. These findings in amphibians and other taxa beg the question of what explains this variation in GC abundance. One idea is that the two GCs are functionally interchangeable, and variation is therefore random. However, given variation even within species, it has been suggested that cortisol and corticosterone may in fact have some different physiological functions and be differentially regulated (Koren et al., 2012; Vera et al., 2019), as suggested by our observations in *D. tinctorius*. The two GCs could be playing different roles across age classes, as suggested by our findings that – using the same sampling methods – corticosterone is seven times as abundant as cortisol in tadpoles (Surber-Cunningham et al., 2024), but the two GCs are only slightly different in adults of *D. tinctorius*. A non-mutually exclusive alternative is that the two GCs could have distinct metabolic functions. For instance, in a study on zebra finches (*Taeniopygia guttata*), the more abundant GC varied by tissue type with more corticosterone (the widely-assumed dominant GC of birds) detected in plasma, but more cortisol detected in immune tissues (Schmidt and Soma, 2008). Additionally, corticosterone and cortisol may differentially bind to species-specific corticosteroid-binding globulins resulting in differing amounts of bound and unbound corticosteroid available in circulation (Breuner and Orchinik, 2002; Martin and Ozon, 1975; Westphal, 1986). Pharmacological studies also suggest differential glucocorticoid receptor-mediated gene regulation based on which type of corticosteroid is available (e.g. Nixon et al., 2016; Pariante et al., 2003) and differential transport of each corticosteroid across the blood-brain barrier (e.g. Karssen et al., 2001). In sum, focusing only on the ACTH-responsive or “dominant” GC ignores ACTH-independent regulation or extra-adrenal production of the other GC, which may be just as physiologically important (Bornstein et al., 2008; Bornstein and Chrousos, 1999; Taves et al., 2011).

### 4.3 Caveats, considerations, and future directions

We end by discussing several caveats and considerations that provide interesting areas for future study. First, we note that many assumptions about GC abundance and dominance are based primarily on studies of plasma or adrenocortical tissue. It is possible that these assumptions do not translate to waterborne hormone sampling. For example, cortisol and corticosterone release rates could differ across species such that waterborne levels of GCs are not representative of circulating levels of GCs in all species. We think it is unlikely that mechanism leading to a difference in release rates would be different among closely related species, thus the differences we see among species likely reflect actual physiological species differences. Additionally, there is precedent for our findings as previous studies examining ACTH responsiveness and relative abundance of GC across many different sample types in amphibians -including water, fecal, saliva, and skin swabs - have demonstrated variation in GC dominance across species. In any case, our results provide evidence that the Dendrobatid species we tested produce substantial amounts of cortisol, contrary to assumptions that amphibians only produce corticosterone.

Second, waterborne hormones are an integrative measure of hormone release rate over time, rather than measuring one moment in time as in plasma. Thus, the collection of waterborne hormones and plasma is inherently separated in time and therefore, perhaps our null expectation should not be a tight relationship between plasma and waterborne hormones. For instance, the duration of waterborne hormone collection may impact the strength of the correlation between plasma and water (e.g., shorter collection timeframe may be more correlated with plasma).

Alternatively, even when plasma and water hormone levels are correlated, GCs may differ in their release rates such that relative abundances differ between plasma and water. For example, in túngara frogs (*Physalaemus pustulosus*), cortisol was present in higher concentrations than corticosterone in waterborne samples, but corticosterone was higher in plasma samples (Baugh et al., 2018). Alternative non-lethal methods for measuring hormones, such as collecting saliva (Hammond et al., 2020, 2018; Tornabene et al., 2023), may be better correlated with plasma hormones given both techniques give a snapshot of hormone concentrations, yet this remains to be tested. In any case, even when correlations between waterborne and plasma samples are imperfect, waterborne samples provide a non-invasive method for measuring biologically meaningful differences in glucocorticoids in anurans (Ruthsatz et al., 2023; Tornabene et al., 2021) and understanding the complexities of this relationship provides interesting opportunities for further study.

Finally, there are inherent limitations with non-lethal hormone sampling. For example, hormones are generally less abundant in water samples than plasma samples and thus can be more difficult to measure. Limited abundance of hormones also makes re-measurement of samples challenging as the concentration can only be diluted so much while remaining within detectable limits of the assay. We had several water samples that did not measure in the range of our ELISA standards. Most of these samples were for our smaller *Ranitomeya* species and we chose to conservatively include them as censored values. While the exact concentration of hormone was not known, the fact that we could confidently determine whether the value was too high or too low to quantify by our curve still provided valuable information. We note that corticosterone, but not cortisol, was particularly difficult to measure in samples from *R. variabilis*, likely due to much lower concentrations of excreted corticosterone than cortisol.

Additionally, while excluding the censored values did reduce our sample sizes, it did not change the overall patterns we observed suggesting there is a robust trend for higher cortisol than corticosterone in multiple species of poison frogs. Despite the limitations of non-invasive methods, these methods are critical in species of conservation concern and small-bodied species where non-lethal techniques are necessary for studies on GC physiology.

### 4.5 Conclusions

Altogether, our findings highlight the importance of measuring both cortisol and corticosterone when investigating the complex, multivariate roles of GCs in physiology, development, and behavior. These findings inspire more questions than they answer. For instance, why does relative GC abundance vary across poison frogs? Are cortisol and corticosterone independently regulated through ACTH and non-ACTH routes? Are poison frogs unique among terrestrial frogs in producing more cortisol than corticosterone? We have a lot left to learn about the stress physiology of this unique clade of frogs and amphibians in general. Based on our findings across Dendrobatid species, we can say with confidence that there is interesting, under-appreciated, and unexplored variation in HPA activity and glucocorticoids.

We encourage more researchers to investigate both GCs in their own focal species. Even if only one GC increases in response to a stressor, the other GC can still play an important role, with non-ACTH sensitive responses providing opportunities for modular regulation of physiology and behavior. Even in well-studies species, measuring both GCs can bring to light new information about the differential regulation and functionality of cortisol and corticosterone, which may not be as interchangeable as we assume (Hancock, 2010). With data across more species, we can look for broader patterns in GC dominance and abundance to investigate whether there are adaptive and/or ecological explanations for the diverse patterns emerging across species, populations, and life stages. We can also test hypotheses about potential unique physiological roles for cortisol versus corticosterone. Additionally, measuring both cortisol and corticosterone may better inform conservation practices through improving our knowledge about stress physiology across species and environments. By understanding the physiological stress processes, we can better consider how to improve conservation efforts to minimize potential stress-related fitness consequences which could impact the species’ survival (Busch and Hayward, 2009; Hofer and East, 1998).

## CRediT authorship contribution statement

**SEW:** conceptualization, methodology, validation, formal analysis, investigation, data curation, writing – original draft, writing – review & editing, visualization, project administration, funding acquisition. **RTP:** conceptualization, methodology, writing – review & editing, supervision. **EKF:** conceptualization, methodology, writing – review & editing, supervision, funding acquisition.

## Acknowledgements

We thank members of the Fischer Lab for their feedback on the manuscript and for the caretaking of the frogs.

## Data Availability Statement

The data underlying this article are available on FigShare, at https://doi.org/10.6084/m9.figshare.23537409.v2.

